# Weighting of sensory cues reflect changing patterns of visual investment during ecological divergence in *Heliconius* butterflies

**DOI:** 10.1101/2024.04.03.587949

**Authors:** José Borrero, Elisa Mogollon Perez, Daniel Shane Wright, Daniela Lozano, Geraldine Rueda-Muñoz, Carolina Pardo-Diaz, Camilo Salazar, Stephen H. Montgomery, Richard M. Merrill

## Abstract

Integrating information across sensory modalities enables animals to orchestrate a wide range of complex behaviours. The relative importance placed on one sensory modality over another reflects the reliability of cues in a particular environment and corresponding differences in neural investment. As populations diverge across environmental gradients, the reliability of sensory cues may shift, favouring divergence in neural investment and the weight given to different sensory modalities. During their divergence across closed-forest and forest-edge habitats, closely related butterflies *Heliconius cydno* and *H. melpomene* evolved distinct brain morphologies, with the former investing more in vision. Quantitative genetic analyses suggest selection drove these changes, but their behavioural effects remain uncertain. We hypothesised that divergent neural investment may alter sensory weighting. We trained individuals in an associative learning experiment using multimodal colour and odour cues. When positively rewarded stimuli were presented in conflict pairing positively trained colour with negatively trained odour, and *vice-versa*, *H. cydno* favoured visual cues more strongly than *H. melpomene*. Hence, differences in sensory weighting may evolve early during divergence and are predicted by patterns of neural investment. These findings, alongside other examples, imply that differences in sensory weighting stem from divergent investment as adaptations to local sensory environments.

## Introduction

Animals rely on multiple sensory systems to perceive and navigate their environments. The integration of environmental information across different sensory modalities (e.g., vision and olfaction) facilitates complex behaviours such as foraging, predator avoidance, and mate finding [1,2]. Cues perceived by these different sensory modalities can work in concert to enhance signal detection [3], but individuals may also prioritise one sensory modality over another [1,4]. This prioritisation of different sensory modalities is likely shaped by divergent selection acting on heritable variation across different habitats [5,6], leading to evolutionary shifts in their relative importance between populations that exploit different habitats.

Evidence for divergence in sensory weighting, defined here as the relative emphasis given to different modalities during behavioural decision-making [4,7,8], has been observed across a handful of taxa where populations or closely related species exploit different sensory environments. For instance, stingless bees rely less on visual cues, such as colour, than honeybees, possibly due to reduced investment in eye size [9]. In butterflies and moths, brain areas processing sensory information are among the most variable structure between species, and studies investigating multisensory integration emphasise associations between interspecific differences in neuroanatomy and behaviour [10,11]. For example, while hawkmoths broadly use both visual and olfactory cues during foraging [4,12], nocturnal species invest relatively more in brain regions associated with olfactory processing, whereas diurnal species are characterised by larger investments in visual neuropils. This shift in neural investment is mirrored in the relative importance given to olfactory and visual cues during foraging [7]. How such specialisations in the weighting of sensory cues evolve over smaller evolutionary time scales and across more subtle ecological gradients at the early stages of divergence is less clear.

The Neotropical *Heliconius* butterflies provide an excellent opportunity to study how sensory environments can shape multisensory integration during the early stages of divergence. Although olfaction also plays a role [8,13–17], these butterflies rely heavily on vision during foraging [18,19], mate recognition [20–22] and oviposition [23,24]. In *Heliconius*, speciation is often associated with ecological transitions across forest types [25–28], such that closely related species experience different sensory conditions [26,29,30]. Sister taxa are also known to differ in multiple aspects of both the peripheral and central sensory pathways, likely reflecting adaptation to local sensory conditions [31–33].

If these shifts in neural investment and sensory weighting are the result of divergent selection imposed by differences in sensory ecology, we expect populations diverging across similar environmental gradients to show associated changes in behaviour [34–36]. The *Heliconius* species pair, *H. melpomene* and *H. cydno,* are a well-studied case of ecological speciation which provide an opportunity to test this hypothesis. These species exist in “mosaic sympatry”, where *H. melpomene* inhabits open forest edges and *H. cydno* occupies more closed canopy, inner forest habitats (Figure 1) [26,37,38]. *H. cydno* has larger eyes and invests more heavily in brain structures associated with vision when compared to *H. melpomene*. Multiple sources of data strongly suggesting these differences have likely been shaped by adaptive processes [32,33,39]. In *Heliconius*, visual information is relayed from the optic lobes to the mushroom bodies (the major sensory integration centres), which also receives input from the olfactory pathway [11]. Changes in investment in specific sensory pathways may therefore affect the strength of input received, consequently influencing the weight given to different sensory cues during behavioural decision-making processes.

**Figure 1:**
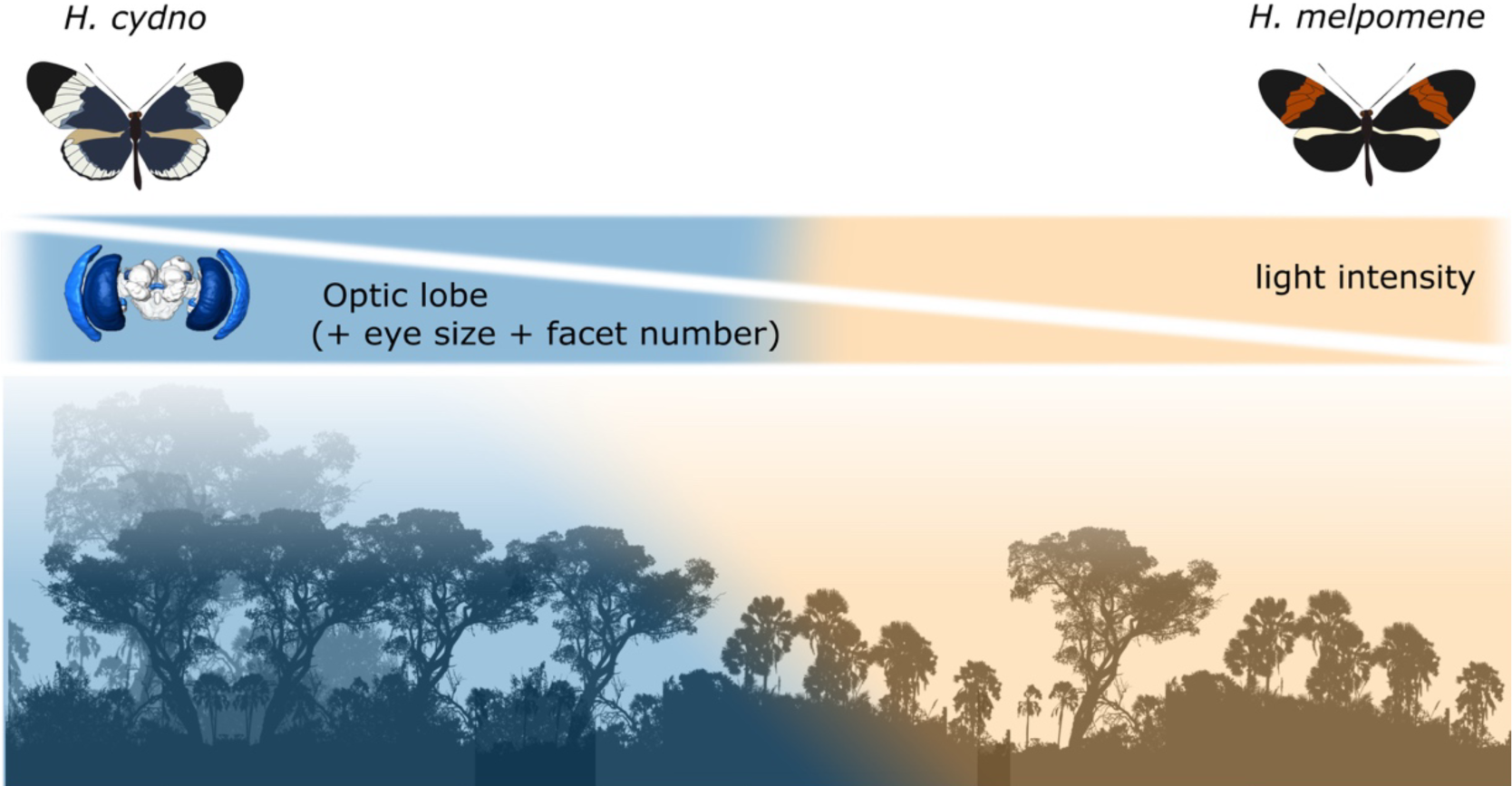
Habitat preference and the sensory environment are associated with differences in brain investment in *Heliconius* butterflies. *H. cydno* invests more in visual neuropils [32] visual periphery [33, 39] and inhabits more-closed canopy forest habitats, than its close relative, *H. melpomene*, which is found in the forest edge [21, 30].

Here, we test how *H. cydno* and *H. melpomene* prioritise visual and olfactory information during foraging, predicting that increased investment in the visual pathway may lead to *H. cydno* prioritising visual cues. To this end, we trained butterflies to a combination of visual and olfactory stimuli using flower models, and then quantified the relative weights given to visual and olfactory cues by presenting the rewarded stimuli in conflict.

## Materials & Methods

### Butterfly rearing and maintenance

We maintained outbred stock populations of *H. cydno* and *H. melpomene* at the Experimental Station José Celestino Mutis – Universidad del Rosario, near La Vega, Colombia (5.0005° N, 74.3394° W), established from wild individuals caught in the local area. Outdoor cages (1×3×2 m) housed butterfly groups with access to a ∼20% sugar solution*, Gurania, Lantana* and *Psiguria spp.* flowers as pollen sources, and *Passiflora* vines for oviposition. All individuals used in the experiments were reared under common garden conditions.

### Experimental set-up and stimuli

To test for visual and olfactory preferences, learning and sensory weighting, we presented male and female butterflies with artificial feeders with both visual (red or blue) and olfactory (lavender or rose essential oils) stimuli on two 25×25 cm hard PVC sheet ‘arrays’ separated by 200 cm in an 1×3×2m insectary (Figure 2A). Each array had four feeders made from a 5 ml Eppendorf tube surrounded by a 3 cm radius circle printed on waterproof paper. One array all red circles, and on the other all blue. Olfactory stimuli were presented in a 200µl PCR-tube below the paper circles, containing either lavender (*Lavandula angustifolia*) or rose *(Rosa damascena*) essential oils diluted with paraffin oil to 1% concentration (Figure 2B). By keeping the olfactory cues separate underneath the waterproof paper and not mixing with the feeding solution, we ensured that olfactory associations were not confounded by gustatory reception. All four feeders on the same array had the same olfactory stimuli to avoid odour mixing.

**Figure 2:**
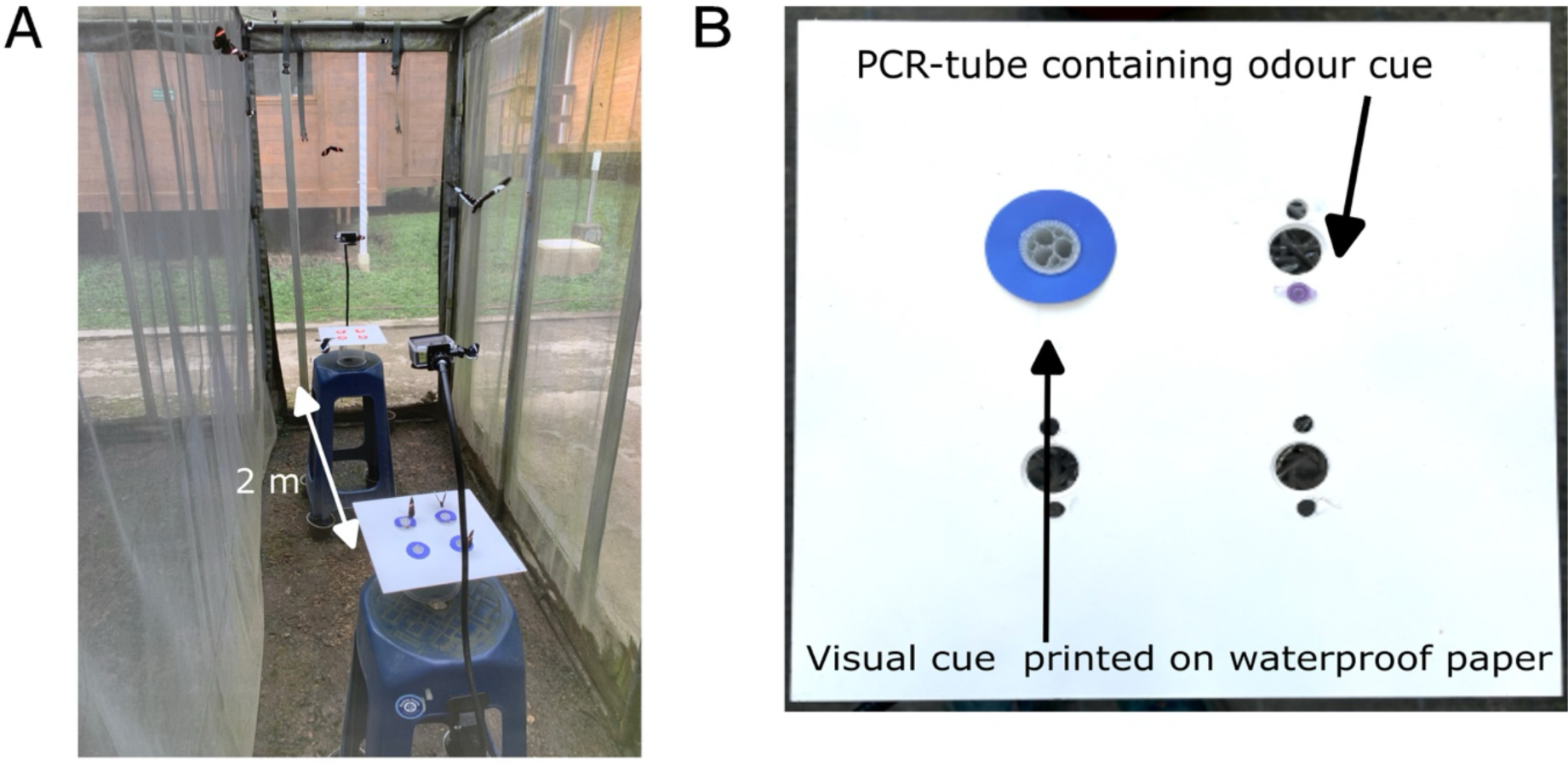
Experimental set-up. **A)** Groups of *H. cydno* and *H. melpomene* were presented with artificial flower arrays at the end of the cages and were trained to a certain colour/odour combination. **B)** Artificial flower array containing the multimodal visual cues (coloured circles) and olfactory cue (PCR tube containing essential oil).

We trained and tested butterflies in same sex groups of 20-24 uniquely marked individuals (10-12 per species). Butterflies were introduced to the experimental cages at least 12 hours prior to the initial testing period to allow for acclimatisation and were food deprived overnight. Butterflies ware trained to associate the combination of blue colour and rose odour with a positive reward [blue/rose] (20% sugar-water solution) and the combination of [red/lavender] with a negative stimulus (∼18% Vitamin C solution; [8,40]). Butterflies were tested with the visual and olfactory in the *trained* test (i.e. [blue/rose] vs [red/lavender]: days 1 and 5), in ‘*conflict’* (i.e. [blue/lavender] vs [red/rose]: day 6), or with *colour alone* (i.e. [blue] vs [red]: day 7) (Figure 3A). In contrast to the training phases, during testing, clean Eppendorf tubes were only filled with water (i.e., with no positive or negative ‘reward’) and fresh visual stimuli were used.

**Figure 3:**
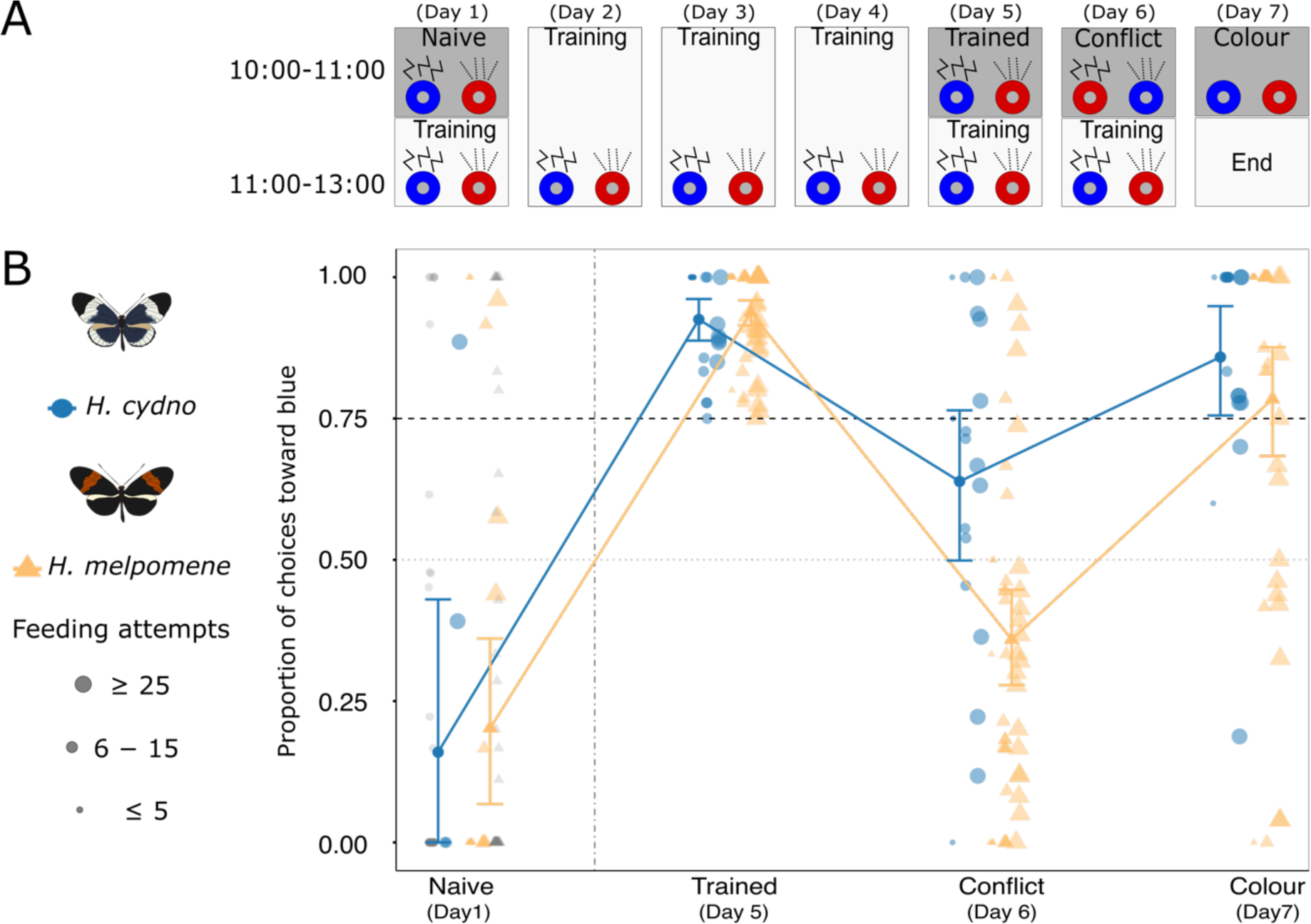
*H. cydno* and *H. melpomene* weight visual and olfactory cues differently while foraging. **A)** Butterflies were trained to feed on blue feeders with a rose oil odour and negatively reinforced red feeders with a lavender smell. In the conflict test, the positively trained feeders were presented in combination with the negatively trained odours and vice versa. During training positive stimuli were rewarded with sugar water and negative stimuli with (∼18% Vitamin C), but during all tests (naive, trained, conflict and colour) feeders were only filled with water (i.e., with no positive or negative reward). Training and reinforcement training periods are delineated by white rectangles, while behavioural tests are indicated by grey rectangles (water only). **B)** P**r**oportion of feeding attempts towards blue feeder for *H. cydno* and *H. melpomene*. Each point represents the feeding choices of an individual; solid dots and error bars represent the mean choices per species ±95% confidence intervals. Blue circles represent *H. cydno* and orange triangles represent *H. melpomene*, shapes in grey represent naive individuals that did not train. On naive and trained tests, feeder combinations were blue/rose and red/lavender; in the conflict test, the positively trained colour (blue) was presented in conflict with the negatively rewarded odour (lavender) and *vice versa*. In the colour tests, individuals were presented only with red or blue feeders without an odour. Only individuals that passed our training criteria (>=75% on the trained day) are shown (see methods).

### Training and testing

i. *Naive test (Day 1)*: We recorded the naive preferences toward combinations of the visual and olfactory stimuli [blue/rose] vs [red/lavender] for one hour (10:00-11:00) using only water feeders. Each butterfly was then hand fed once on a positive [blue/rose (+)], and once on a negative rewarded feeder [red/lavender (-)], and then allowed to feed freely for 2 hours from the positive and negative rewarded feeders (11:00-13:00).
ii. Training (Days 2-4): Training was reinforced on experimental days 2-4. First, each individual was hand fed once on a positive [blue/rose (+)], and once on a negative rewarded feeder [red/lavender (-)]. Afterwards, butterflies were allowed to feed freely for 3 hours (10:00-13:00) from the positive or negative rewarded feeders. The feeders’ position was randomised (coin toss) every day to prevent positional learning.
iii. *Trained test* (Day 5): On day 5, we recorded the trained preferences for one hour (between 10:00-11:00) using only water in the feeders with the stimuli in *trained* conditions (i.e. [blue/rose] vs [red/lavender]). This was immediately followed by two hours of reinforcement training with rewarded feeders, i.e. [blue/rose (+)] and [red/lavender (-)].
iv. *Conflict test* (Day 6): To test for differences in sensory weighting, we presented the visual and olfactory in conflict (i.e. [blue/lavender] vs [red/rose]; Figure 3A). As for the previous tests, feeding behaviour was recorded for one hour (between 10:00-11:00) using only water in the feeders. This was then followed by reinforcement training for two hours, i.e. [blue/rose (+)] and [red/lavender (-)].
v. *Colour test* (Day 7): We tested visual preferences in isolation for one hour (between 10:00-11:00) (i.e., blue vs red, without olfactory stimuli, and using water only).

### Behavioural tests

All feeding test assays [(i) Naive, (ii) Trained, (iii) Conflict and (iv) Colour] were filmed from above with an AKASO Brave 4 Pro action camera. We scored the videos in BORIS [41], recording the total number of feeding attempts directed toward each stimulus per individual. Feeding attempts were defined as a proboscis extension within the coloured circle feeders. We scored instances where an individual moved outside the circle and subsequently returned as a new feeding attempt. In contrast, we scored instances where an individual remained within a circle and probed continuously as a single attempt, unless the proboscis was retracted for 3 seconds or longer.

### Training criterion

We established a stringent training criterion to incorporate only individuals that had successfully learned the correct stimulus combination, following similar learning studies in lizards [42,43] and butterflies [19]. Out of 260 individuals, 73 were excluded from the analysis for not attempting to feed on the trained test (day 5) (Table S2). To investigate species differences in sensory weighting, we included only those individuals that demonstrated robust associative learning of the trained cue combinations (i.e. >=75% of feeding choices towards the positive multimodal stimuli [blue/rose] on the trained test (day 5)), resulting in a reduced sample size from 187 to 73 individuals. Although biased towards *H. melpomene* (52 individuals vs 21 *H. cydno* individuals), this rigorous criterion ensured that our analysis focused solely on individuals who had successfully learned. However, our results did not quantitatively change when including all 117 individuals with a lower threshold of learning (i.e. >50% correct; see supplementary results Table S1 and Tables S5-S6).

### Statistical analysis

We used general linear mixed models (GLMM) with a binomial distribution in the *lme4* [44] in R [45] to assess if species differed in their feeding attempts when the positively rewarded stimuli agreed or were in conflict. The dependent variable was the proportion of choices toward the blue feeders and we included species, and treatment [Trained test (day 5), Conflict test (day 6) or Colour test (day 7)], and their interaction as fixed factors. To account for the varying number of choices made by each butterfly, individual observations are weighted by their total number of feeding attempts [46]. Individual ID nested within group ID was included as random effects to account for repeated measures (Table S1). We applied a stepwise model reduction approach to identify the minimum adequate statistical models, using likelihood ratio tests (LRT) via the *drop1* function to determine significance. When fixed effects or their interactions were significant, we conducted pairwise comparisons tests using post hoc Bonferroni corrections with the *emmeans* package [47]. Plots were made using *ggplot2* [48].

## Results

Among the 73 individuals (52 *H. melpomene* and 21 *H. cydno*) that passed our stringent training criterion (i.e., >75% correct choices on day 5), we conducted tests to quantify the species’ reliance on visual and olfactory cues. Importantly, during these tests, all feeders were unrewarded. *H. cydno* did not differ from *H. melpomene* in the trained test (Table 1 post-hoc: *H. cydno* Trained vs. *H. melpomene* Trained, Z= −1.375, P= 1.0). In contrast, *H. melpomene* and *H. cydno* responded differently when presented with the olfactory and visual stimuli in conflict, (Figure 3B). Importantly, the interaction between species and trial type was retained in our model, confirming that this was an effect of how the stimuli were presented (species*treatment LRT: 2ΔlnL = 22.876, df=2, p < 0.001. Table S3). In the conflict test, *H. cydno* fed mostly on the blue feeder, while *H. melpomene* primarily chose the feeder with the rose scent, suggesting a difference between these species in how olfactory and visual cues are weighed during behavioural decisions (Table 1, post-hoc: *H. cydno* Conflict vs. *H. melpomene* Conflict, Z= 3.210, P= 0.019). We found no effect of sex, which was excluded from our model (sex LRT: 2ΔlnL = 3.067, df=1, p = 0.079. Table S3).

**Table 1:** Pairwise p-value comparison matrix of post-hoc test with Bonferroni correction for the minimal adequate model for the effects of the interaction between species and treatment (Trained, Conflict, and Colour) on choices towards blue feeders. Only individuals that passed our training criteria (>=75% on the trained day) are included Results are from a binomial GLMM: *see methods for statistical details. Proportion blue ∼ Species * Level + (1|Group/ID)*

|  | *H. cydno*<br>Trained | *H. melpomene*<br>Trained | *H. cydno*<br>Conflict | *H. melpomene*<br>Conflict | *H. cydno*<br>Colour | *H. melpomene*<br>Colour |
| --- | --- | --- | --- | --- | --- | --- |
| *H. cydno* Trained | [ 2.329] | 1 | <.0001 | <.0001 | 0.3041 | 0.008 |
| *H. melpomene*<br>Trained |  | [ 2.874] | <.0001 | <.0001 | 0.0116 | <.0001 |
| *H. cydno* Conflict |  |  | [ 0.465] | 0.0199 | <.0001 | 1 |
| *H. melpomene*<br>Conflict |  |  |  | [-0.601] | <.0001 | <.0001 |
| *H. cydno* Colour |  |  |  |  | [1.639] | 1 |
| *H. melpomene*<br>Colour |  |  |  |  |  | [0.989] |

Our post-hoc analysis revealed that individuals of both species made fewer feeding attempts on the blue feeder when the stimuli were presented in conflict, suggesting that both species use both visual and olfactory stimuli when making foraging decisions (Table 1, post-hoc: *H. cydno* Trained vs. *H. cydno* Conflict, Z= 6.932, P= <.0001; *H. melpomene* Trained vs. *H. melpomene* Conflict, Z=20.614, P= <.0001). *H. cydno* made fewer feeding attempts on the blue feeder when the positively trained olfactory and visual stimuli were in conflict, than when they were in the trained combinations confirming the use of both cues when present. However, there was no distinction between the trained test and the colour-only test (Table 1, post-hoc: *H. cydno* Trained vs. *H. cydno* Colour, Z= 2.321, P= 0.304), further emphasising that *H. cydno* was more attentive to the visual cue in our assay. Conversely, *H. melpomene* differed between the trained test and the colour-only test, indicating greater use of the olfactory stimulus (Table 1, post-hoc: *H. melpomene* Trained vs. *H. melpomene* Colour, Z= 10.920, P= <.0001)

## Discussion

We explored how two closely related butterfly species, *H. cydno* and *H. melpomene*, weigh visual and olfactory information in a foraging context. Our findings indicate a significant divergence in behaviour between these two species, with *H. cydno* favouring visual cues over olfactory cues compared to *H. melpomene.* Importantly, our results mirror differences in brain morphology and suggest that divergence in the sensory environment may have led to changes in brain composition and how these species process, integrate, and prioritise information across sensory modalities.

To assess how *H. cydno* and *H. melpomene* prioritise visual and olfactory information, we presented trained butterflies with conflicting stimuli - pairing positively trained colours with negatively trained odours, and *vice versa*. Both species chose the ‘correct’ colour less frequently in these conflict scenarios compared to their trained preferences or when presented with colours alone, indicating that both visual and olfactory cues influence foraging decisions. However, when presented with conflicting cues, the species differed in their reliance on each sensory modality*. H. cydno* predominantly fed on the positively rewarded visual cue (the blue feeder). In contrast, *H. melpomene* showed a stronger preference for the feeder containing the positive trained odour stimulus (rose), indicating a difference between two species in the weight given to the two cues during behavioural decision-making. Additionally, *H. melpomene* differed between the trained and colour only test, making fewer correct choices when only colours were presented, again suggesting a larger reliance on the olfaction stimulus compared to *H. cydno*. As the visual and olfactory cues employed in our study are not assumed to be ecologically relevant, we suggest they reflect the baseline relative preferences of the species for each sensory modality. Given that the behavioural variation we observe is from common-garden-reared individuals, these interspecific shifts are likely to have a genetic basis. In addition, our results align with divergent patterns of investment in sensory pathways between the two species, whereby *H. cydno* exhibits greater relative investment in the visual system [32,33,39]. Therefore, heritable shifts in neural investment and sensory weighting may result from divergent selection imposed by differences in sensory ecology.

One potential limitation of our study is that we managed to successfully train a larger number of *H. melpomene* than *H. cydno* individuals. This discrepancy in training could be due to *H. melpomene* being more active during the trials, possibly due to the outdoor insectaries being found in the forest edge, which closely resembles the *H. melpomene* habitat [26,30]. Indeed, in contrast to *H. cydno, H. melpomene* is frequently observed flying next to the insectaries (pers. obs. J. Borrero). Despite this difference, individuals of both species successfully learned the rewarded combination under our rigorous training criteria (>= 75% correct choices) and our main conclusions hold even under less rigorous training criteria (see supplementary results).

Adaptation to the local sensory environment may play an important role during the early stages of divergence. The environmental gradient observed in our studied species, *H. cydno* and *H. melpomene*, from closed forest canopy to open forest edge [26,30], resembles the transition across an altitudinal gradient, from complex broad-leaf forest environments to dry Andean forests seen in another distantly related *Heliconius* species-pair, *H. erato* and *H. himera* [49]. *H. erato cyrbia* and *H. himera* exhibit similar patterns of neural divergence, with *H. erato cyrbia* showing increased, heritable investment in the visual pathway, but with *H. himera* also showing some increased investment in the olfactory pathway [8,31]. Despite methodological differences (group vs individual testing and the exact visual and olfactory stimuli used), our results qualitatively reassemble a previous study in *H. erato* and *H. himera* [8]. Together, these findings demonstrate similar shifts in sensory weighting among *Heliconius* butterflies, with species inhabiting more closed habitats investing more in perceiving and processing visual information and favouring visual over olfactory cues. This suggests changes in the light environment is a common source of divergent selection across the two species pairs.

More broadly, changes in sensory weighting have been observed across different taxa and across different evolutionary timescales. For instance, urban Great tits prefer olfactory cues, while forest birds rely more on visual cues [50]. Nine-spined sticklebacks show plasticity in their sensory reliance, for instance compared to fish reared in visually unrestricted habitats, individuals raised in visually restricted environments display a larger reliance on chemical over visual cues. This suggests neural plasticity may underpin initial behavioural responses to novel environments that are later consolidated by selection [51]. In threespine stickleback fish, populations in different lakes segregate into limnetic and benthic ecomorphs along an environmental gradient [52]. These stickleback populations show repeated evolutionary shifts in brain regions that are correlated with differences in their sensory environments; limnetic species have larger structures for visual processing and benthic ectomorphs possess larger olfactory bulbs. Comparing across larger evolutionary timescales, studies have found that nocturnal hawkmoths prioritise olfactory cues while foraging, as evidenced by their increased investment in olfactory processing. Conversely, diurnal hawkmoths show larger investment in visual neuropils and place a greater emphasis on visual cues. These changes in the relative weight given to different sensory modalities, and the accompanying changes in brain investment, are likely driven by the transition from a nocturnal to a diurnal habitat and the associated alterations in their sensory environment [7]. Combined with our findings, these studies underscore the role of environmental factors in driving changes in neuroanatomy, sensory integration, and behaviour across species.

In conclusion, our results highlight shifts in sensory weighting between species associated with neural investment and ecological divergence, consistent with patterns across *Heliconius* butterflies. We demonstrated that *H. cydno*, from a lower light habitat [30] which is characterised with increased visual investment [32,33], favours visual cues more strongly than *H. melpomene*. Alongside previous results in *Heliconius* butterflies and other taxa making [7,8], our research reveals repeated shifts in sensory weighting associated with changes in neural investment and habitat use. This highlights the potential role for the sensory environment to shape differences in brain structure, which subsequently may influence key sensory behaviours crucial during the initial stages of species divergence.

## Acknowledgements

We thank Universidad del Rosario for insectary access at JCM and Isabel León for assistance, as well as Amulya Hosur and Alejandro Mancino for video scoring help.

## Supplementary material

### Supplementary Results

Among the 117 individuals (82 *H. melpomene* and 35 *H. cydno*) that passed our less stringent training criterion (i.e., >50% correct choices on day 5), *H. cydno* did not differ from *H. melpomene* in the trained test, indicating that both species were able to train to the multimodal stimulus (Table S1 post-hoc: *H. cydno* Trained vs. *H. melpomene* Trained, Z= −1.333, P= 1.0). However, *H. melpomene* and *H. cydno* responded differently when presented with the olfactory and visual stimuli in conflict (Figure S1). Importantly, the interaction between species and trial type was retained in our model, confirming that this was not due to existing differences amongst “trained” individuals, but rather the effect of how the stimuli were presented (species*treatment LRT: 2ΔlnL = 44.537, df=2, p < 0.001). Table S5). In the conflict test, *H. cydno* fed mostly on the blue feeder, while *H. melpomene* primarily chose the feeder with the rose scent, suggesting a difference between these species in how olfactory and visual cues are weighed during behavioural decisions (Table S1, post-hoc: *H. cydno* Conflict vs. *H. melpomene* Conflict, Z= 4.402, P< 0.001). We found no effect of sex, which was excluded from our model (sex LRT: 2ΔlnL = 0.297, df=1, p = 0.586. Table S5).

Post-hoc analysis revealed that individuals of both species made fewer feeding attempts on the blue feeder when the stimuli were presented in conflict, indicating that both species use both visual and olfactory stimuli when making foraging decisions (Table S1, post-hoc: *H. cydno* Trained vs. *H. cydno* Conflict, Z= 5.760, P= <.0001) (Table S1, post-hoc: *H. melpomene* Trained vs. *H. melpomene* Conflict, Z=23.222, P= <.0001). Notably, *H. cydno* did not differ from the trained test and the colour-only test indicating that this species largely relies on visual cues (Table S1, post-hoc: *H. cydno* Trained vs. *H. cydno* Conflict, Z= 0.337, P= 1.0). In contrast, *H. melpomene* differed between the trained test and the colour-only test further suggesting a difference in sensory weighting (Table 1, post-hoc: *H. melpomene* Trained vs. *H. melpomene* Conflict, Z= 8.433, P= <.0001).

**Figure S 1:**
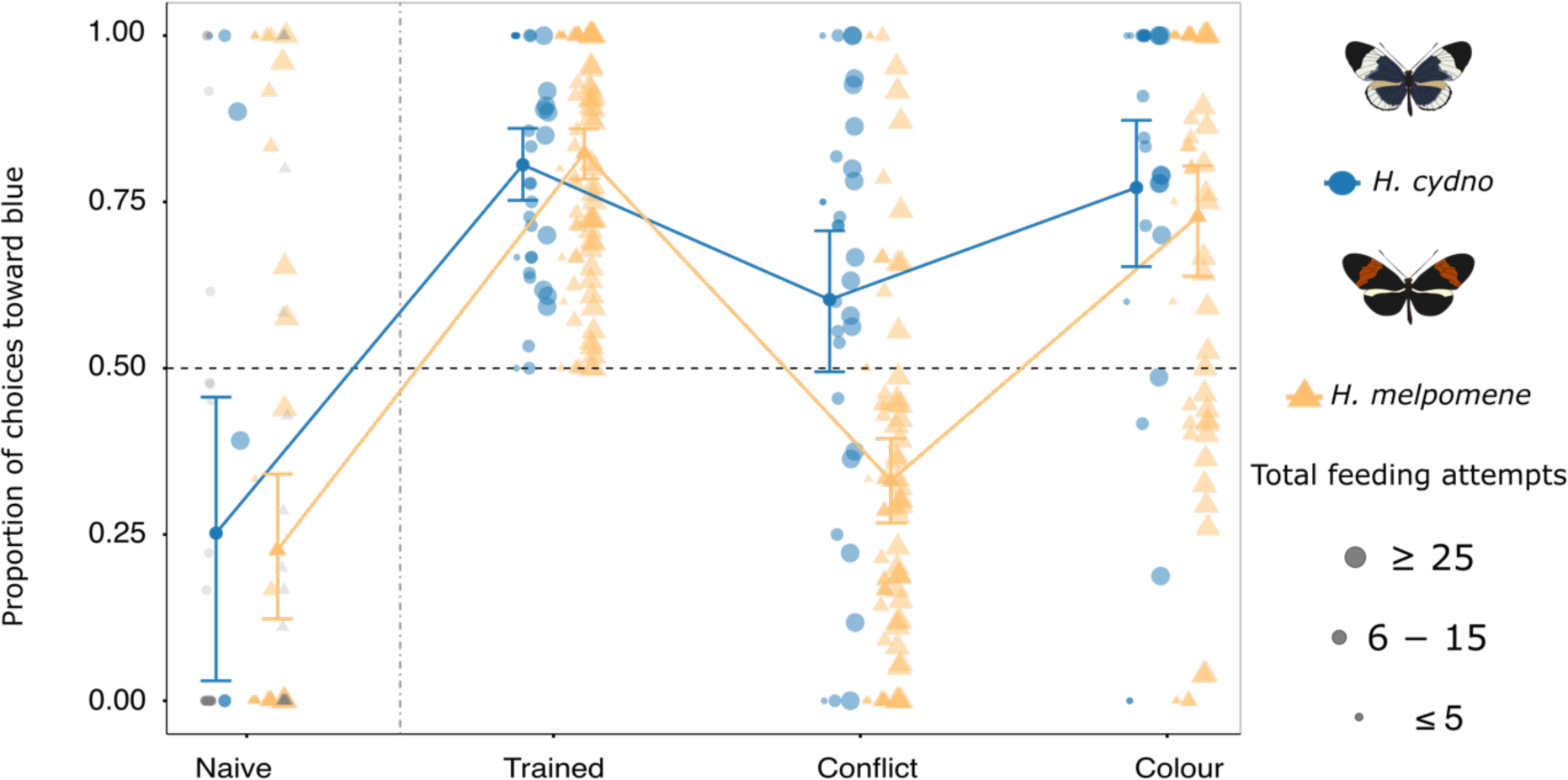
Our result that *H. cydno* and *H. melpomene* weight visual and olfactory cues differently when foraging is also supported when adopting a less stringent learning criteria. Proportion of feeding attempts towards blue feeder for *H. cydno* and *H. melpomene*. Each point represents the feeding choices of and individual; solid dots and error bars represent the mean choices per species ±95% confidence intervals. Blue circles represent *H. cydno* and orange triangles represent *H. melpomene*, shapes in grey represent naive individuals that did not train. On naive and trained tests, feeder combinations were blue/rose and red/lavender; In the conflict test, the positively trained colour (blue) was presented in conflict with the negatively rewarded odour (lavender) and *vice versa*. In the colour tests, individuals were presented only with red or blue feeders without an odour. Only individuals that passed our training criteria (>=50% on the trained day) are show (see methods).

**Table S 1:** Pairwise p-value comparison matrix of post-hoc test with Bonferroni correction for the minimal adequate model for the effects of the interaction between species, treatment (Trained, Conflict, and Colour) and their interaction on choices towards blue feeders. Only individuals that passed our training criteria (>=50% on the trained day) are on this dataset. Results are from binomial GLMM model, see methods for statistical details.

|  | *H. cydno*<br>Trained | *H. melpomene*<br>Trained | *H. cydno*<br>Conflict | *H. melpomene*<br>Conflict | *H. cydno*<br>Colour | *H. melpomene*<br>Colour |
| --- | --- | --- | --- | --- | --- | --- |
| *H. cydno* Trained | [1.331] | 1 | <.0001 | <.0001 | 1 | 0.338 |
| *H. melpomene* Trained |  | [1.656] | <.0001 | <.0001 | 1 | <.0001 |
| *H. cydno* Conflict |  |  | [0.360] | 0.0002 | <.0001 | 1 |
| *H. melpomene* Conflict |  |  |  | [-0.649] | <.0001 | <.0001 |
| *H. cydno* Colour |  |  |  |  | [1.265] | 0.7534 |
| *H. melpomene* Colour |  |  |  |  |  | [0.771] |

**Figure S2:**
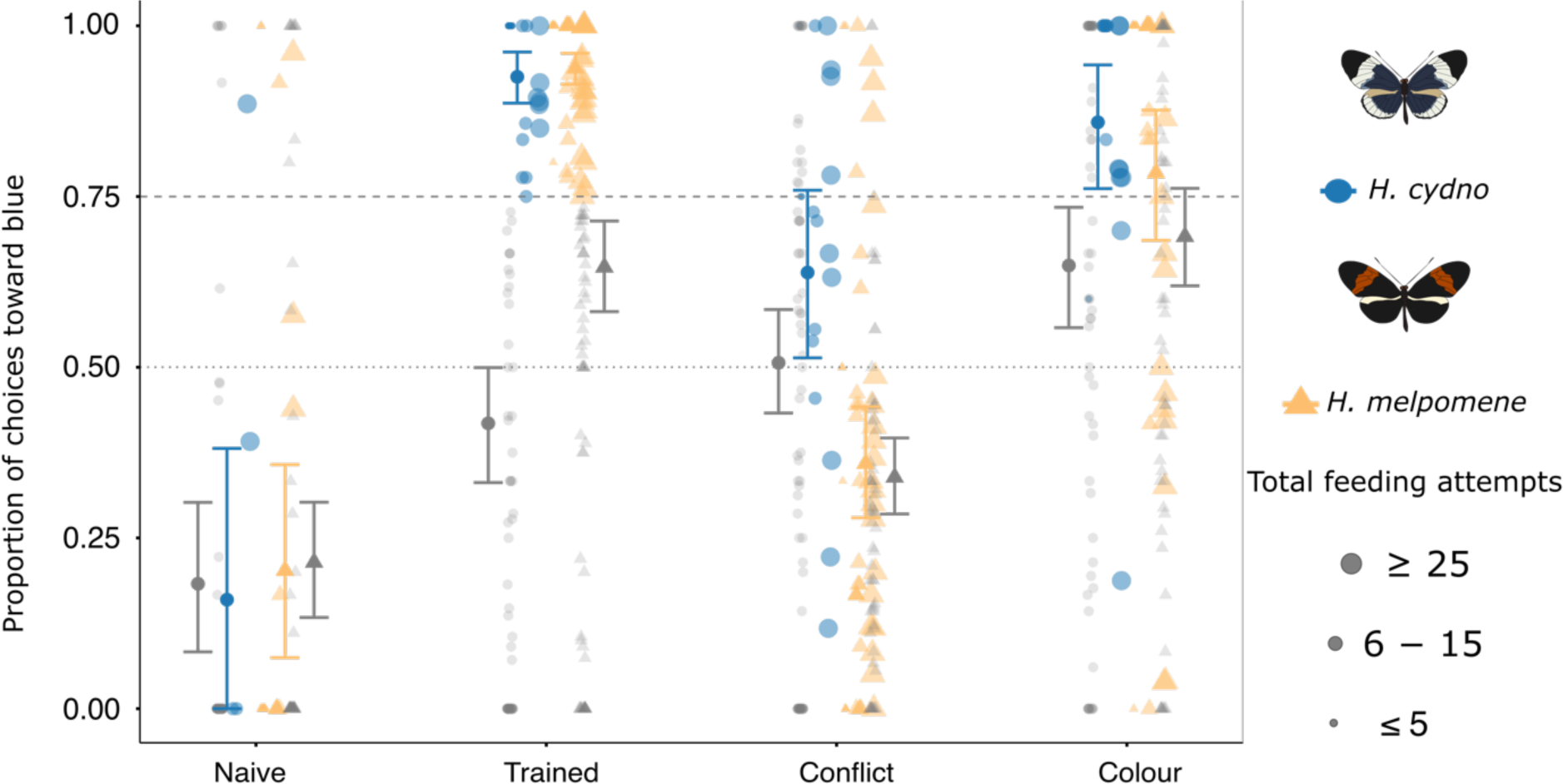
Behavioural choices including individuals that did not reach our training criterion (>=75%). Proportion of feeding attempts towards blue feeder for *H. cydno* and *H. melpomene*. Each point represents the feeding choices of and individual; solid dots and error bars represent the mean choices per species ±95% confidence intervals. Blue circles represent *H. cydno* and orange triangles represent *H. melpomene*, shapes in grey represent individuals that did not train. Gray error bars represent the mean choices per species ±95% confidence intervals including all individuals.

**Table S 2:** Number of individuals by sex that participated in each treatment (Naïve, Trained, Conflict, Colour) separated by number of individuals that passed the training criterion (>=50% or >=75%) of correct choices on the trained day (day 5).

| Species | Level | Sex | All individuals | $\geq 50\%$ training | $\geq 75\%$ training |
| --- | --- | --- | --- | --- | --- |
| *H. cydno* | Naive | Female | 23 | 5 | 2 |
| *H. cydno* | Naive | Male | 38 | 8 | 6 |
| *H. cydno* | Trained | Female | 30 | 13 | 6 |
| *H. cydno* | Trained | Male | 49 | 22 | 15 |
| *H. cydno* | Conflict | Female | 35 | 11 | 5 |
| *H. cydno* | Conflict | Male | 56 | 20 | 14 |
| *H. cydno* | Colour | Female | 29 | 9 | 6 |
| *H. cydno* | Colour | Male | 48 | 18 | 12 |
| *H. melpomene* | Naive | Female | 23 | 14 | 10 |
| *H. melpomene* | Naive | Male | 51 | 34 | 15 |
| *H. melpomene* | Trained | Female | 40 | 30 | 24 |
| *H. melpomene* | Trained | Male | 68 | 52 | 28 |
| *H. melpomene* | Conflict | Female | 41 | 26 | 20 |
| *H. melpomene* | Conflict | Male | 68 | 48 | 28 |
| *H. melpomene* | Colour | Female | 35 | 21 | 16 |
| *H. melpomene* | Colour | Male | 61 | 45 | 26 |

**Figure S3:**
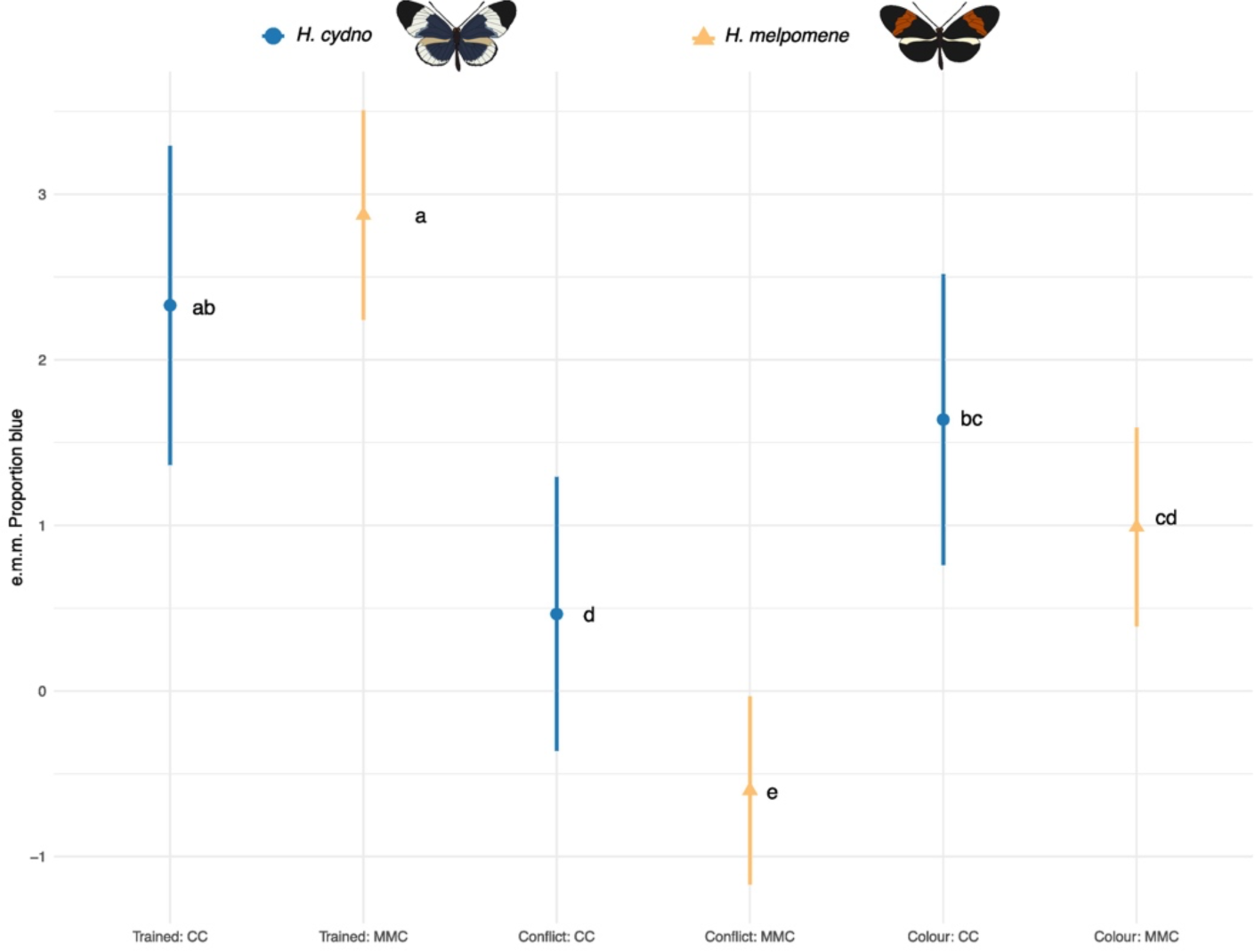
Estimated marginal means (e.m.m.) of the minimum adequate statistical models for the proportion of feeding attempts towards blue feeder demonstrating the significant effects of the interaction between species (*H. cydno* and *H. melpomene*) and treatment [Trained test (day 5), Conflict test (day 6) or Colour test (day 7)]. Only individuals that passed our training criteria (>=75% on the trained day) are on this dataset. (see methods).

**Table S 3:** Model 1 minimal adequate model for the effects of species, treatment (Trained, Conflict, Colour) and their interaction on choices towards blue feeders. Only individuals that passed our training criteria (>=75% on the trained day) are on this dataset. Results are from a binomial GLMM model, see methods for statistical details.

|  | 2ΔlnL | df | Pr |
| --- | --- | --- | --- |
| Species | 27.846 | 3 | 3.912e-06 *** |
| Treatment | 681.740 | 4 | < 2.2e-16 *** |
| Treatment:Species | 22.876 | 2 | 1.078e-05 *** |
| Sex | 3.067 | 1 | 0.07988 |

**Table S 4:** Pairwise comparison post-hoc test with Bonferroni correction for the minimal adequate model for the effects of the interaction between species and treatment (Trained, Conflict, and Colour) on choices towards blue feeders. Only individuals that passed our training criteria (>=75% on the trained day) are on this dataset. Results are from binomial GLMM model, see methods for statistical details. Proportion blue ∼ Species * Level + (1|Group/ID)

|  | estimate | SE | z-ratio | p-value<br>(Adjusted) |
| --- | --- | --- | --- | --- |
| *H. cydno* Trained - *H. melpomene* Trained | -0.545 | 0.396 | -1.375 | 1.000 |
| *H. cydno* Trained - *H. cydno* Conflict | 1.864 | 0.269 | 6.932 | <.0001 |
| *H. cydno* Trained - *H. melpomene* Conflict | 2.930 | 0.381 | 7.696 | <.0001 |
| *H. cydno* Trained - *H. cydno* Colour | 0.690 | 0.297 | 2.321 | 0.304 |
| *H. cydno* Trained - *H. melpomene* Colour | 1.340 | 0.387 | 3.464 | 0.008 |
| *H. melpomene* Trained - *H. cydno* Conflict | 2.409 | 0.351 | 6.860 | <.0001 |
| *H. melpomene* Trained - *H. melpomene* Conflict | 3.475 | 0.169 | 20.614 | <.0001 |
| *H. melpomene* Trained - *H. cydno* Colour | 1.235 | 0.367 | 3.361 | 0.012 |
| *H. melpomene* Trained - *H. melpomene* Colour | 1.884 | 0.173 | 10.920 | <.0001 |
| *H. cydno* Conflict - *H. melpomene* Conflict | 1.066 | 0.332 | 3.210 | 0.020 |
| *H. cydno* Conflict - *H. cydno* Colour | -1.174 | 0.237 | -4.956 | <.0001 |
| *H. cydno* Conflict - *H. melpomene* Colour | -0.524 | 0.340 | -1.544 | 1.000 |
| *H. melpomene* Conflict - *H. cydno* Colour | -2.240 | 0.351 | -6.385 | <.0001 |
| *H. melpomene* Conflict - *H. melpomene* Colour | -1.590 | 0.141 | -11.244 | <.0001 |
| *H. cydno* Colour - *H. melpomene* Colour | 0.650 | 0.357 | 1.819 | 1.000 |

**Table S 5:**
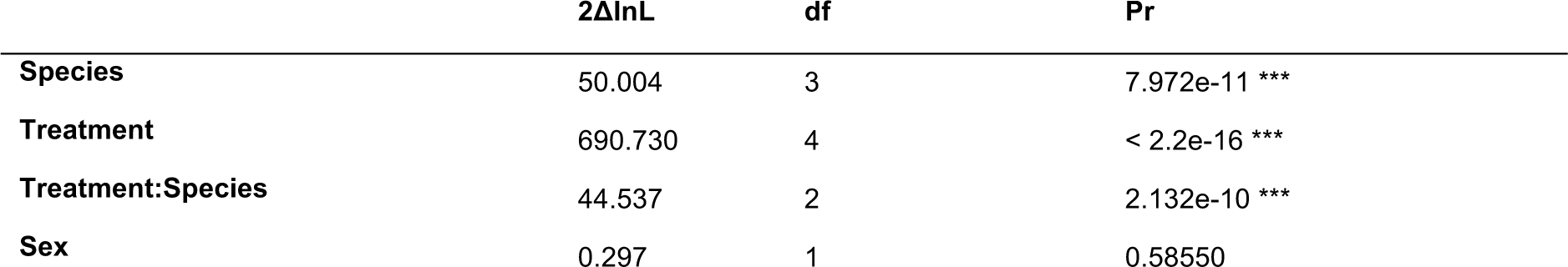
Model 2 minimal adequate model for the effects of species, treatment (Trained, Conflict, Colour) and their interaction on choices towards blue feeders. (>=50% on the trained day) are on this dataset. Results are from binomial GLMM model, see methods for statistical details.

**Figure S4:**
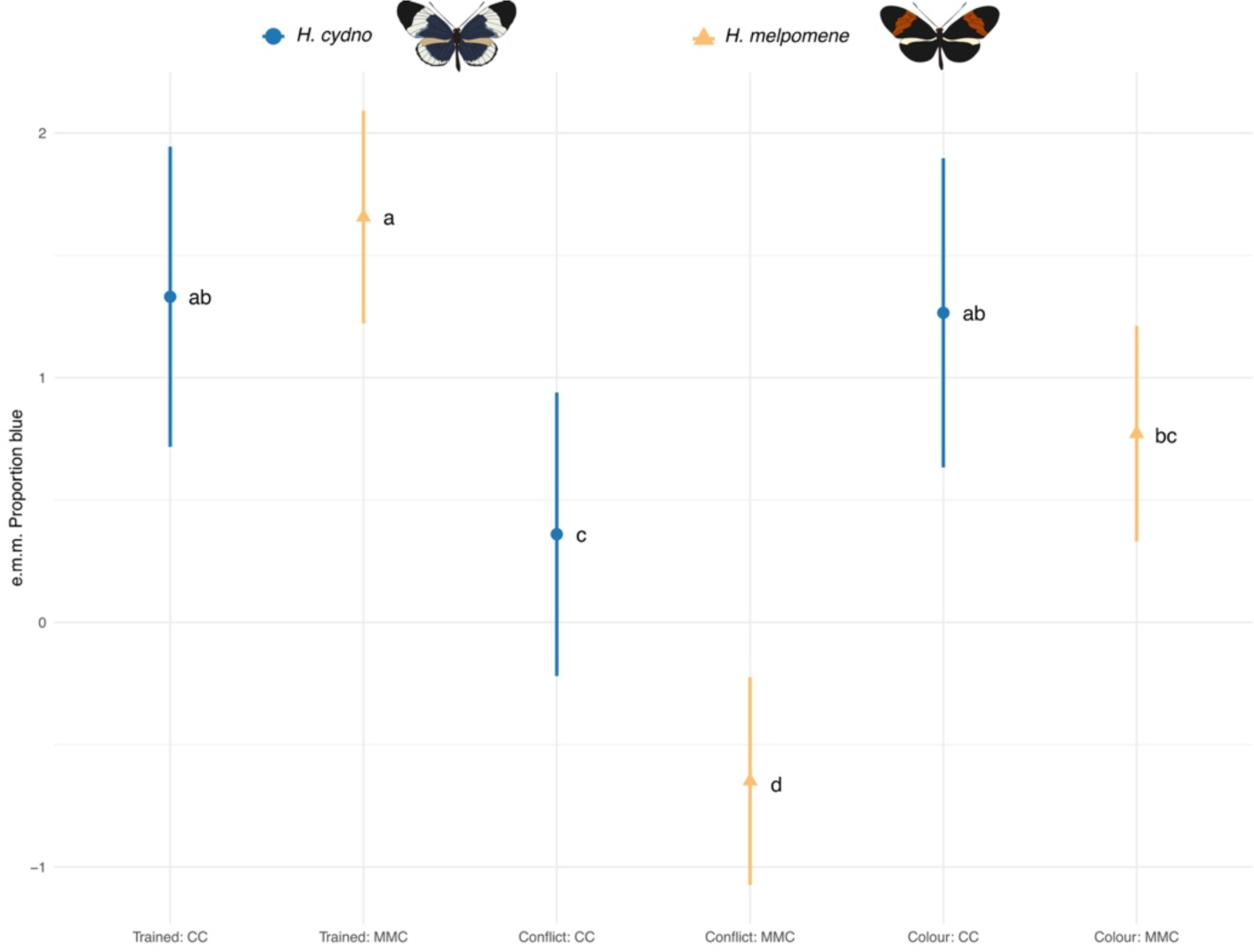
Estimated marginal means (e.m.m.) of the minimum adequate statistical models for the proportion of feeding attempts towards blue feeder demonstrating the significant effects of the interaction between species (*H. cydno* and *H. melpomene)* and treatment [*Trained test* (day 5), *Conflict test* (day 6) or *Colour test* (day 7)]. Only individuals that passed our training criteria (>=50% on the trained day) are on this dataset. (see methods).

**Table S 6:** Pairwise comparison post-hoc test with Bonferroni correction for the minimal adequate model for the effects of the interaction between species and treatment (Trained, Conflict, and Colour) on choices towards blue feeders. Only individuals that passed our training criteria (>=50% on the trained day) are on this dataset. Results are from binomial GLMM model, see methods for statistical details.

|  | estimate | SE | z-ratio | p-value |
| --- | --- | --- | --- | --- |
| *H. cydno* Trained - *H. melpomene* Trained | -0.326 | 0.244 | -1.333 | 1.000 |
| *H. cydno* Trained - *H. cydno* Conflict | 0.971 | 0.169 | 5.760 | <.0001 |
| *H. cydno* Trained - *H. melpomene* Conflict | 1.980 | 0.241 | 8.199 | <.0001 |
| *H. cydno* Trained - *H. cydno* Colour | 0.066 | 0.195 | 0.337 | 1.000 |
| *H. cydno* Trained - *H. melpomene* Colour | 0.560 | 0.245 | 2.281 | 0.338 |
| *H. melpomene* Trained - *H. cydno* Conflict | 1.296 | 0.233 | 5.571 | <.0001 |
| *H. melpomene* Trained - *H. melpomene* Conflict | 2.305 | 0.099 | 23.222 | <.0001 |
| *H. melpomene* Trained - *H. cydno* Colour | 0.391 | 0.251 | 1.556 | 1.000 |
| *H. melpomene* Trained - *H. melpomene* Colour | 0.885 | 0.105 | 8.433 | <.0001 |
| *H. cydno* Conflict - *H. melpomene* Conflict | 1.009 | 0.229 | 4.402 | 0.000 |
| *H. cydno* Conflict - *H. cydno* Colour | -0.905 | 0.183 | -4.942 | <.0001 |
| *H. cydno* Conflict - *H. melpomene* Colour | -0.411 | 0.234 | -1.760 | 1.000 |
| *H. melpomene* Conflict - *H. cydno* Colour | -1.914 | 0.249 | -7.697 | <.0001 |
| *H. melpomene* Conflict - *H. melpomene* Colour | -1.420 | 0.100 | -14.263 | <.0001 |
| *H. cydno* Colour - *H. melpomene* Colour | 0.494 | 0.252 | 1.958 | 0.753 |

**Table S7:**
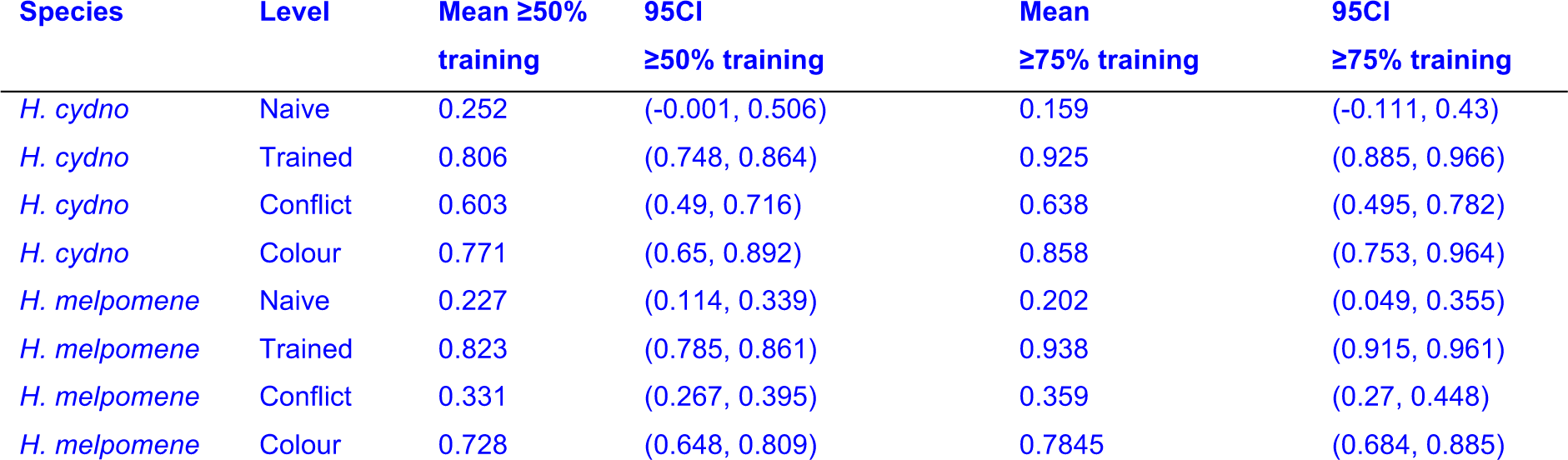
Average proportion of feeding attempts separated by species and treatment (Naïve, Trained, Conflict, Colour) that passed the training criterion (>=50% or >=75%) of correct choices on the trained day (day 5).

**Table S 8:**
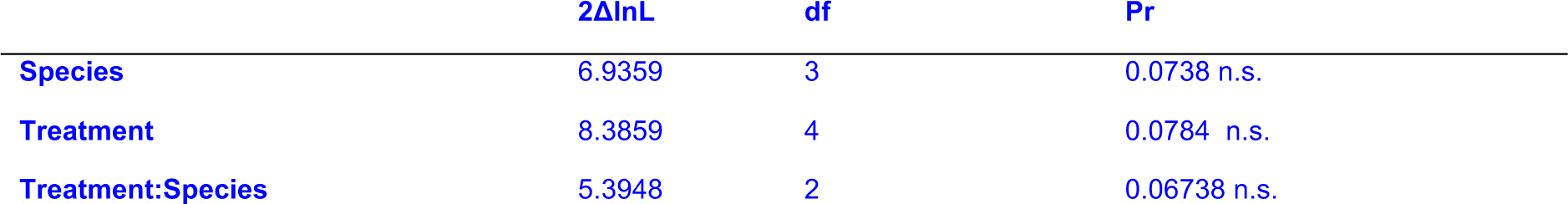
Model X minimal adequate model for the effects of species, treatment (Trained, Conflict, Colour) and their interaction on the total number of feeding attempts. Only individuals that passed our training criteria (>=75% on the trained day 5) are on this dataset. Results are from a binomial GLMM model, see methods for statistical details. Total_feeding_attempts ∼ Species * Level + (1|Group/ID)

**Table S9:** Total number of feeding attempts and proportion of feeding attempts towards blue feeders grouped by individual. Only individuals that passed our training criteria (>=75% on the trained day5) are on this dataset.

| ID | Species | Sex | Level | Day | Red | Blue | Total | Prop_blue | Group |
| --- | --- | --- | --- | --- | --- | --- | --- | --- | --- |
| C22_004 | MMC | M | Naive | 1 | 0 | 3 | 3 | 1 | G14 |
| C22_107 | CC | F | Naive | 1 | 1 | 0 | 1 | 0 | G15 |
| C22_091 | MMC | M | Naive | 1 | 2 | 0 | 2 | 0 | G16 |
| C22_173 | MMC | F | Naive | 1 | 0 | 1 | 1 | 1 | G17 |
| C22_226 | MMC | F | Naive | 1 | 3 | 0 | 3 | 0 | G17 |
| C22_273 | MMC | F | Naive | 1 | 2 | 0 | 2 | 0 | G17 |
| C22_028 | MMC | M | Naive | 1 | 9 | 0 | 9 | 0 | G18 |
| C22_433 | MMC | F | Naive | 1 | 1 | 0 | 1 | 0 | G20 |
| C22_590 | MMC | F | Naive | 1 | 1 | 0 | 1 | 0 | G21 |
| C22_831 | MMC | F | Naive | 1 | 1 | 0 | 1 | 0 | G21 |
| C22_898 | MMC | F | Naive | 1 | 12 | 0 | 12 | 0 | G21 |
| C22_918 | MMC | F | Naive | 1 | 8 | 0 | 8 | 0 | G21 |
| C22_555 | MMC | M | Naive | 1 | 7 | 0 | 7 | 0 | G22 |
| C22_699 | CC | M | Naive | 1 | 4 | 31 | 35 | 0 | G22 |
| A23_055 | MMC | M | Naive | 1 | 23 | 18 | 41 | 0 | G22 |
| A23_094 | MMC | M | Naive | 1 | 1 | 0 | 1 | 0 | G23 |
| C22_704 | MMC | M | Naive | 1 | 14 | 19 | 33 | 0 | G23 |
| A23_280 | CC | F | Naive | 1 | 1 | 0 | 1 | 0 | G25 |
| A23_054 | CC | M | Naive | 1 | 14 | 9 | 23 | 0 | G25 |
| A23_242 | MMC | M | Naive | 1 | 5 | 1 | 6 | 0 | G25 |
| A23_306 | MMC | M | Naive | 1 | 1 | 11 | 12 | 0 | G25 |
| C22_911 | MMC | M | Naive | 1 | 7 | 0 | 7 | 0 | G26 |
| C22_935 | CC | M | Naive | 1 | 1 | 0 | 1 | 0 | G26 |
| C22_936 | CC | M | Naive | 1 | 6 | 0 | 6 | 0 | G26 |
| A23_098 | MMC | M | Naive | 1 | 1 | 0 | 1 | 0 | G27 |
| A23_148 | MMC | M | Naive | 1 | 1 | 0 | 1 | 0 | G27 |
| A23_182 | MMC | M | Naive | 1 | 1 | 0 | 1 | 0 | G27 |
| A23_258 | MMC | M | Naive | 1 | 1 | 24 | 25 | 0 | G25 |
| A23_259 | MMC | M | Naive | 1 | 3 | 0 | 3 | 0 | G27 |
| A23_303 | CC | M | Naive | 1 | 5 | 0 | 5 | 0 | G27 |
| A23_428 | CC | M | Naive | 1 | 6 | 0 | 6 | 0 | G27 |
| A23_224 | MMC | F | Naive | 1 | 2 | 0 | 2 | 0 | G28 |
| A23_283 | MMC | F | Naive | 1 | 1 | 0 | 1 | 0 | G28 |
| C22_004 | MMC | M | Trained | 5 | 0 | 16 | 16 | 1 | G14 |
| C22_063 | MMC | F | Trained | 5 | 13 | 52 | 65 | 0 | G25 |
| C22_107 | CC | F | Trained | 5 | 0 | 2 | 2 | 1 | G15 |
| C22_134 | MMC | F | Trained | 5 | 0 | 1 | 1 | 1 | G15 |
| C22_025 | CC | M | Trained | 5 | 0 | 1 | 1 | 1 | G16 |
| C22_032 | MMC | M | Trained | 5 | 0 | 6 | 6 | 1 | G16 |
| C22_091 | MMC | M | Trained | 5 | 0 | 26 | 26 | 1 | G16 |
| C22_173 | MMC | F | Trained | 5 | 0 | 1 | 1 | 1 | G17 |
| C22_198 | CC | F | Trained | 5 | 3 | 17 | 20 | 0 | G17 |
| C22_226 | MMC | F | Trained | 5 | 2 | 22 | 24 | 0 | G17 |
| C22_265 | MMC | F | Trained | 5 | 0 | 20 | 20 | 1 | G17 |
| C22_273 | MMC | F | Trained | 5 | 0 | 9 | 9 | 1 | G17 |
| C22_285 | MMC | F | Trained | 5 | 0 | 49 | 49 | 1 | G17 |
| C22_028 | MMC | M | Trained | 5 | 0 | 1 | 1 | 1 | G18 |
| C22_242 | CC | M | Trained | 5 | 2 | 7 | 9 | 0 | G25 |
| C22_227 | MMC | M | Trained | 5 | 3 | 11 | 14 | 0 | G25 |
| C22_290 | CC | M | Trained | 5 | 0 | 6 | 6 | 1 | G19 |
| C22_341 | MMC | M | Trained | 5 | 12 | 36 | 48 | 0 | G19 |
| C22_367 | MMC | M | Trained | 5 | 2 | 12 | 14 | 0 | G19 |
| C22_448 | MMC | M | Trained | 5 | 4 | 39 | 43 | 0 | G19 |
| C22_186 | MMC | F | Trained | 5 | 1 | 4 | 5 | 0 | G19 |
| C22_230 | MMC | F | Trained | 5 | 2 | 14 | 16 | 0 | G19 |
| C22_422 | MMC | F | Trained | 5 | 2 | 24 | 26 | 0 | G19 |
| C22_433 | MMC | F | Trained | 5 | 0 | 46 | 46 | 1 | G20 |
| C22_590 | MMC | F | Trained | 5 | 0 | 11 | 11 | 1 | G21 |
| C22_831 | MMC | F | Trained | 5 | 0 | 8 | 8 | 1 | G21 |
| C22_898 | MMC | F | Trained | 5 | 2 | 16 | 18 | 0 | G21 |
| C22_918 | MMC | F | Trained | 5 | 0 | 26 | 26 | 1 | G21 |
| C22_555 | MMC | M | Trained | 5 | 0 | 4 | 4 | 1 | G22 |
| C22_699 | CC | M | Trained | 5 | 0 | 5 | 5 | 1 | G22 |
| C22_756 | MMC | M | Trained | 5 | 0 | 1 | 1 | 1 | G22 |
| A23_055 | MMC | M | Trained | 5 | 0 | 4 | 4 | 1 | G23 |
| A23_094 | MMC | M | Trained | 5 | 2 | 7 | 9 | 0 | G23 |
| C22_704 | MMC | M | Trained | 5 | 0 | 7 | 7 | 1 | G23 |
| C22_848 | MMC | M | Trained | 5 | 3 | 20 | 23 | 0 | G23 |
| A23_264 | MMC | F | Trained | 5 | 0 | 3 | 3 | 1 | G25 |
| A23_280 | CC | F | Trained | 5 | 2 | 6 | 8 | 0 | G25 |
| A23_030 | MMC | M | Trained | 5 | 0 | 1 | 1 | 1 | G26 |
| A23_054 | CC | M | Trained | 5 | 2 | 7 | 9 | 0 | G26 |
| A23_185 | MMC | M | Trained | 5 | 0 | 4 | 4 | 1 | G26 |
| A23_242 | MMC | M | Trained | 5 | 1 | 13 | 14 | 0 | G26 |
| A23_301 | CC | M | Trained | 5 | 0 | 7 | 7 | 1 | G26 |
| A23_306 | MMC | M | Trained | 5 | 0 | 3 | 3 | 1 | G26 |
| A23_319 | CC | M | Trained | 5 | 0 | 5 | 5 | 1 | G26 |
| C22_843 | CC | M | Trained | 5 | 0 | 4 | 4 | 1 | G26 |
| C22_911 | MMC | M | Trained | 5 | 5 | 16 | 21 | 0 | G26 |
| C22_935 | CC | M | Trained | 5 | 0 | 1 | 1 | 1 | G26 |
| C22_936 | CC | M | Trained | 5 | 3 | 23 | 26 | 0 | G26 |
| C22_949 | CC | M | Trained | 5 | 0 | 2 | 2 | 1 | G26 |
| A23_056 | MMC | M | Trained | 5 | 5 | 17 | 22 | 0 | G26 |
| A23_083 | CC | M | Trained | 5 | 0 | 18 | 18 | 1 | G27 |
| A23_098 | MMC | M | Trained | 5 | 1 | 19 | 20 | 0 | G27 |
| A23_148 | MMC | M | Trained | 5 | 1 | 21 | 22 | 0 | G27 |
| A23_182 | MMC | M | Trained | 5 | 4 | 33 | 37 | 0 | G27 |
| A23_258 | MMC | M | Trained | 5 | 0 | 2 | 2 | 1 | G27 |
| A23_259 | MMC | M | Trained | 5 | 1 | 5 | 6 | 0 | G27 |
| A23_268 | MMC | F | Trained | 5 | 0 | 11 | 11 | 1 | G27 |
| A23_289 | MMC | M | Trained | 5 | 0 | 19 | 19 | 1 | G27 |
| A23_303 | CC | M | Trained | 5 | 1 | 6 | 7 | 0 | G27 |
| A23_345 | MMC | M | Trained | 5 | 0 | 8 | 8 | 1 | G27 |
| A23_347 | MMC | M | Trained | 5 | 1 | 10 | 11 | 0 | G27 |
| A23_421 | CC | M | Trained | 5 | 2 | 17 | 19 | 0 | G27 |
| A23_428 | CC | M | Trained | 5 | 2 | 16 | 18 | 0 | G27 |
| A23_009 | CC | F | Trained | 5 | 0 | 5 | 5 | 1 | G28 |
| A23_027 | CC | F | Trained | 5 | 1 | 5 | 6 | 0 | G28 |
| A23_209 | CC | F | Trained | 5 | 5 | 55 | 60 | 0 | G28 |
| A23_221 | MMC | F | Trained | 5 | 0 | 10 | 10 | 1 | G28 |
| A23_224 | MMC | F | Trained | 5 | 0 | 33 | 33 | 1 | G28 |
| A23_283 | MMC | F | Trained | 5 | 0 | 4 | 4 | 1 | G28 |
| C22_809 | MMC | F | Trained | 5 | 5 | 21 | 26 | 0 | G28 |
| C22_891 | MMC | F | Trained | 5 | 0 | 8 | 8 | 1 | G28 |
| C22_900 | MMC | F | Trained | 5 | 3 | 28 | 31 | 0 | G28 |
| C22_920 | MMC | F | Trained | 5 | 3 | 28 | 31 | 0 | G28 |
| C22_004 | MMC | M | Conflict | 6 | 1 | 20 | 21 | 0 | G28 |
| C22_063 | MMC | F | Conflict | 6 | 22 | 11 | 33 | 0 | G28 |
| C22_134 | MMC | F | Conflict | 6 | 5 | 0 | 5 | 0 | G15 |
| C22_025 | CC | M | Conflict | 6 | 5 | 0 | 5 | 0 | G16 |
| C22_032 | MMC | M | Conflict | 6 | 2 | 22 | 24 | 0 | G16 |
| C22_091 | MMC | M | Conflict | 6 | 5 | 8 | 13 | 0 | G16 |
| C22_198 | CC | F | Conflict | 6 | 14 | 8 | 22 | 0 | G16 |
| C22_226 | MMC | F | Conflict | 6 | 15 | 3 | 18 | 0 | G16 |
| C22_265 | MMC | F | Conflict | 6 | 39 | 15 | 54 | 0 | G16 |
| C22_273 | MMC | F | Conflict | 6 | 8 | 0 | 8 | 0 | G17 |
| C22_285 | MMC | F | Conflict | 6 | 21 | 9 | 30 | 0 | G17 |
| C22_028 | MMC | M | Conflict | 6 | 30 | 4 | 34 | 0 | G17 |
| C22_242 | CC | M | Conflict | 6 | 0 | 21 | 21 | 1 | G18 |
| C22_227 | MMC | M | Conflict | 6 | 5 | 0 | 5 | 0 | G19 |
| C22_341 | MMC | M | Conflict | 6 | 4 | 2 | 6 | 0 | G19 |
| C22_367 | MMC | M | Conflict | 6 | 19 | 1 | 20 | 0 | G19 |
| C22_448 | MMC | M | Conflict | 6 | 5 | 1 | 6 | 0 | G19 |
| C22_422 | MMC | F | Conflict | 6 | 10 | 1 | 11 | 0 | G19 |
| C22_433 | MMC | F | Conflict | 6 | 10 | 2 | 12 | 0 | G19 |
| C22_590 | MMC | F | Conflict | 6 | 17 | 12 | 29 | 0 | G19 |
| C22_831 | MMC | F | Conflict | 6 | 0 | 3 | 3 | 1 | G21 |
| C22_898 | MMC | F | Conflict | 6 | 34 | 3 | 37 | 0 | G21 |
| C22_555 | MMC | M | Conflict | 6 | 3 | 11 | 14 | 0 | G21 |
| C22_699 | CC | M | Conflict | 6 | 1 | 3 | 4 | 0 | G21 |
| C22_756 | MMC | M | Conflict | 6 | 0 | 13 | 13 | 1 | G22 |
| A23_055 | MMC | M | Conflict | 6 | 5 | 4 | 9 | 0 | G22 |
| A23_094 | MMC | M | Conflict | 6 | 20 | 0 | 20 | 0 | G23 |
| C22_704 | MMC | M | Conflict | 6 | 10 | 28 | 38 | 0 | G23 |
| C22_848 | MMC | M | Conflict | 6 | 11 | 3 | 14 | 0 | G23 |
| A23_264 | MMC | F | Conflict | 6 | 1 | 0 | 1 | 0 | G25 |
| A23_280 | CC | F | Conflict | 6 | 7 | 12 | 19 | 0 | G25 |
| A23_030 | MMC | M | Conflict | 6 | 10 | 2 | 12 | 0 | G25 |
| A23_054 | CC | M | Conflict | 6 | 2 | 25 | 27 | 0 | G25 |
| A23_185 | MMC | M | Conflict | 6 | 5 | 10 | 15 | 0 | G25 |
| A23_242 | MMC | M | Conflict | 6 | 4 | 27 | 31 | 0 | G25 |
| A23_301 | CC | M | Conflict | 6 | 9 | 18 | 27 | 0 | G25 |
| A23_306 | MMC | M | Conflict | 6 | 19 | 18 | 37 | 0 | G25 |
| A23_319 | CC | M | Conflict | 6 | 0 | 8 | 8 | 1 | G26 |
| C22_843 | CC | M | Conflict | 6 | 3 | 8 | 11 | 0 | G26 |
| C22_911 | MMC | M | Conflict | 6 | 31 | 20 | 51 | 0 | G26 |
| C22_935 | CC | M | Conflict | 6 | 2 | 5 | 7 | 0 | G26 |
| C22_936 | CC | M | Conflict | 6 | 7 | 25 | 32 | 0 | G26 |
| C22_949 | CC | M | Conflict | 6 | 4 | 5 | 9 | 0 | G26 |
| A23_056 | MMC | M | Conflict | 6 | 17 | 8 | 25 | 0 | G26 |
| A23_083 | CC | M | Conflict | 6 | 14 | 4 | 18 | 0 | G26 |
| A23_098 | MMC | M | Conflict | 6 | 22 | 3 | 25 | 0 | G26 |
| A23_148 | MMC | M | Conflict | 6 | 9 | 2 | 11 | 0 | G26 |
| A23_182 | MMC | M | Conflict | 6 | 28 | 7 | 35 | 0 | G26 |
| A23_258 | MMC | M | Conflict | 6 | 8 | 6 | 14 | 0 | G26 |
| A23_259 | MMC | M | Conflict | 6 | 2 | 2 | 4 | 0 | G26 |
| A23_268 | MMC | F | Conflict | 6 | 7 | 0 | 7 | 0 | G27 |
| A23_289 | MMC | M | Conflict | 6 | 10 | 8 | 18 | 0 | G27 |
| A23_303 | CC | M | Conflict | 6 | 6 | 7 | 13 | 0 | G27 |
| A23_345 | MMC | M | Conflict | 6 | 2 | 1 | 3 | 0 | G27 |
| A23_347 | MMC | M | Conflict | 6 | 5 | 4 | 9 | 0 | G27 |
| A23_421 | CC | M | Conflict | 6 | 15 | 2 | 17 | 0 | G27 |
| A23_428 | CC | M | Conflict | 6 | 6 | 5 | 11 | 0 | G27 |
| A23_009 | CC | F | Conflict | 6 | 0 | 2 | 2 | 1 | G28 |
| A23_027 | CC | F | Conflict | 6 | 1 | 3 | 4 | 0 | G28 |
| A23_209 | CC | F | Conflict | 6 | 2 | 29 | 31 | 0 | G28 |
| A23_221 | MMC | F | Conflict | 6 | 12 | 0 | 12 | 0 | G28 |
| A23_224 | MMC | F | Conflict | 6 | 7 | 6 | 13 | 0 | G28 |
| A23_283 | MMC | F | Conflict | 6 | 19 | 11 | 30 | 0 | G28 |
| C22_809 | MMC | F | Conflict | 6 | 1 | 1 | 2 | 0 | G28 |
| C22_891 | MMC | F | Conflict | 6 | 0 | 2 | 2 | 1 | G28 |
| C22_900 | MMC | F | Conflict | 6 | 6 | 0 | 6 | 0 | G28 |
| C22_920 | MMC | F | Conflict | 6 | 9 | 2 | 11 | 0 | G28 |
| C22_004 | MMC | M | Colour | Day7 | 10 | 18 | 28 | 0 | G28 |
| C22_063 | MMC | F | Colour | Day7 | 22 | 16 | 38 | 0 | G28 |
| C22_107 | CC | F | Colour | Day7 | 13 | 3 | 16 | 0 | G28 |
| C22_134 | MMC | F | Colour | Day7 | 0 | 14 | 14 | 1 | G15 |
| C22_032 | MMC | M | Colour | Day7 | 0 | 5 | 5 | 1 | G16 |
| C22_091 | MMC | M | Colour | Day7 | 0 | 14 | 14 | 1 | G16 |
| C22_173 | MMC | F | Colour | Day7 | 8 | 0 | 8 | 0 | G17 |
| C22_198 | CC | F | Colour | Day7 | 0 | 17 | 17 | 1 | G17 |
| C22_226 | MMC | F | Colour | Day7 | 0 | 1 | 1 | 1 | G17 |
| C22_265 | MMC | F | Colour | Day7 | 2 | 11 | 13 | 0 | G17 |
| C22_285 | MMC | F | Colour | Day7 | 9 | 7 | 16 | 0 | G17 |
| C22_227 | MMC | M | Colour | Day7 | 7 | 5 | 12 | 0 | G17 |
| C22_290 | CC | M | Colour | Day7 | 4 | 15 | 19 | 0 | G17 |
| C22_341 | MMC | M | Colour | Day7 | 1 | 5 | 6 | 0 | G17 |
| C22_367 | MMC | M | Colour | Day7 | 24 | 1 | 25 | 0 | G17 |
| C22_448 | MMC | M | Colour | Day7 | 7 | 14 | 21 | 0 | G17 |
| C22_590 | MMC | F | Colour | Day7 | 0 | 6 | 6 | 1 | G21 |
| C22_831 | MMC | F | Colour | Day7 | 0 | 4 | 4 | 1 | G21 |
| C22_898 | MMC | F | Colour | Day7 | 25 | 1 | 26 | 0 | G21 |
| C22_555 | MMC | M | Colour | Day7 | 3 | 19 | 22 | 0 | G21 |
| C22_699 | CC | M | Colour | Day7 | 2 | 3 | 5 | 0 | G21 |
| C22_756 | MMC | M | Colour | Day7 | 3 | 0 | 3 | 0 | G22 |
| A23_055 | MMC | M | Colour | Day7 | 0 | 7 | 7 | 1 | G23 |
| A23_094 | MMC | M | Colour | Day7 | 0 | 7 | 7 | 1 | G23 |
| C22_704 | MMC | M | Colour | Day7 | 0 | 9 | 9 | 1 | G23 |
| C22_848 | MMC | M | Colour | Day7 | 0 | 7 | 7 | 1 | G23 |
| A23_280 | CC | F | Colour | Day7 | 4 | 14 | 18 | 0 | G23 |
| A23_030 | MMC | M | Colour | Day7 | 10 | 10 | 20 | 0 | G23 |
| A23_054 | CC | M | Colour | Day7 | 0 | 29 | 29 | 1 | G26 |
| A23_185 | MMC | M | Colour | Day7 | 4 | 12 | 16 | 0 | G26 |
| A23_242 | MMC | M | Colour | Day7 | 0 | 47 | 47 | 1 | G26 |
| A23_301 | CC | M | Colour | Day7 | 0 | 14 | 14 | 1 | G26 |
| A23_306 | MMC | M | Colour | Day7 | 21 | 18 | 39 | 0 | G26 |
| A23_319 | CC | M | Colour | Day7 | 0 | 10 | 10 | 1 | G26 |
| C22_843 | CC | M | Colour | Day7 | 0 | 9 | 9 | 1 | G26 |
| C22_911 | MMC | M | Colour | Day7 | 27 | 13 | 40 | 0 | G26 |
| C22_935 | CC | M | Colour | Day7 | 0 | 6 | 6 | 1 | G26 |
| C22_936 | CC | M | Colour | Day7 | 9 | 21 | 30 | 0 | G26 |
| C22_949 | CC | M | Colour | Day7 | 0 | 7 | 7 | 1 | G26 |
| A23_056 | MMC | M | Colour | Day7 | 0 | 10 | 10 | 1 | G27 |
| A23_083 | CC | M | Colour | Day7 | 0 | 10 | 10 | 1 | G27 |
| A23_098 | MMC | M | Colour | Day7 | 0 | 20 | 20 | 1 | G27 |
| A23_148 | MMC | M | Colour | Day7 | 0 | 3 | 3 | 1 | G27 |
| A23_182 | MMC | M | Colour | Day7 | 0 | 10 | 10 | 1 | G27 |
| A23_258 | MMC | M | Colour | Day7 | 0 | 5 | 5 | 1 | G27 |
| A23_259 | MMC | M | Colour | Day7 | 0 | 8 | 8 | 1 | G27 |
| A23_268 | MMC | F | Colour | Day7 | 0 | 2 | 2 | 1 | G27 |
| A23_289 | MMC | M | Colour | Day7 | 0 | 14 | 14 | 1 | G27 |
| A23_303 | CC | M | Colour | Day7 | 0 | 4 | 4 | 1 | G27 |
| A23_345 | MMC | M | Colour | Day7 | 0 | 3 | 3 | 1 | G27 |
| A23_421 | CC | M | Colour | Day7 | 0 | 6 | 6 | 1 | G27 |
| A23_009 | CC | F | Colour | Day7 | 1 | 5 | 6 | 0 | G27 |
| A23_027 | CC | F | Colour | Day7 | 4 | 14 | 18 | 0 | G27 |
| A23_209 | CC | F | Colour | Day7 | 9 | 34 | 43 | 0 | G27 |
| A23_221 | MMC | F | Colour | Day7 | 0 | 3 | 3 | 1 | G28 |
| A23_224 | MMC | F | Colour | Day7 | 2 | 10 | 12 | 0 | G28 |
| A23_283 | MMC | F | Colour | Day7 | 0 | 5 | 5 | 1 | G28 |
| C22_809 | MMC | F | Colour | Day7 | 1 | 7 | 8 | 0 | G28 |
| C22_900 | MMC | F | Colour | Day7 | 0 | 5 | 5 | 1 | G28 |
| C22_920 | MMC | F | Colour | Day7 | 0 | 5 | 5 | 1 | G28 |

